# Embryogenesis in myrmicine ants combines features of short and long germ-band modes of development

**DOI:** 10.1101/2023.09.19.557849

**Authors:** Chi-Chun Fang, Arjuna Rajakumar, Andrew Kenny, Ulrich G. Mueller, Ehab Abouheif, David Stein

## Abstract

Ants exhibit complex social organization, morphologically distinct castes with division of labor, and the exploitation of diverse ecological niches. The extent to which these features have influenced embryonic development relative to other insects remains unclear. Insect embryogenesis has been classified into one of three modes: long, short, and intermediate germ-band. In long germ-band development, exemplified by the fruit fly *Drosophila melanogaster*, segments along the entire anterior-posterior axis of the embryonic primordium are established almost simultaneously, prior to gastrulation, with the initial embryonic primordium surrounding almost the entire volume of the egg. In short and intermediate germ-band modes, the embryonic primordium occupies a smaller proportion of the egg surface, with anterior segments initially specified, and remaining segments being added sequentially from a posterior growth zone. Here, we show a novel pattern of development in three myrmicine ants, the fungus-gardening ants *Atta texana* and *Mycocepurus smithii*, and the red imported fire ant *Solenopsis invicta*. Early in embryogenesis, they exhibit features of short germ-band development, while later in development they exhibit a newly-characterized progressive pattern of segmentation that has been associated with some long germ-band-developing insects. Moreover, despite similarities in the size of ant and *Drosophila* eggs, the duration of embryogenesis in the three ant species is 10 to 20-fold longer than in *Drosophila* and is also significantly longer than in the honeybee *Apis mellifera* and the jewel wasp *Nasonia vitripennis*. In addition, the embryos produced by *A. texana* foundress queens develop to first instar larvae 25% faster than embryos produced by mature queens. We discuss these results in the context of the eusocial lifestyle of ants.

## Introduction

The ants (Formicidae, Hymenoptera) are a remarkably successful group of insects owing to their social behaviors, diverse morphologies, division of labor among distinct castes, specialized life-history traits, and ecological abundance (Hölldobler & Wilson, 1990; Bourke & Franks, 1995; Kipyatkov & Lopatina, 2015). Despite extensive studies on morphology, behavior, and communication in ants, embryonic development has been understudied. Ant embryogenesis has received some recent attention (Abouheif & Wray, 2002, Khila & Abouheif, 2008, 2010; Niculita, 2006; Rafiki et al., 2020), yet the degree to which patterns of segmentation in ants is conserved relative to other insects remains unknown.

Historically, insect embryogenesis has been classified into three morphologically distinguishable types: long, intermediate, and short germ-band (Krause, 1939; Sander, 1976; Tear et al. 1990; Akam, 1994; Davis & Patel, 2002; Liu & Kaufman, 2005; Lynch & Roth, 2011; Lynch et al., 2012). In embryos undergoing long germ-band development, almost all the cells of the blastoderm contribute to the embryonic primordium such that it fills the entire space within the eggshell and surrounds a central yolk mass. In long germ-band-developing insects, the blastoderm represents the rudiments of all the segments making up the head, thorax, and abdomen of the embryo, with the segmented body plan being generated by subdivision of the initial embryonic primordium. In the best-understood long germ-band-developing insect *Drosophila melanogaster*, studies of early development indicated that specification of all segments occurs almost simultaneously (Akam, 1987; Ingham, 1988), and prior to the onset of gastrulation. At the molecular level, this manifests itself in the almost simultaneous appearance of seven stripes of mRNA encoding each of the members of the pair-rule class of genes, which includes the genes *even-skipped* (*eve*), *hairy* (*h*), *runt* (*run*) and *fushi tarazu* (*ftz*)(Macdonald et al.,1986; Ingham et al.,1985; Gergen and Butler, 1988; Hafen et al.,1984), followed soon after by the appearance of fourteen stripes of the segment polarity genes *wingless* (*wg*), *engrailed* (*en*), and *hedgehog* (*hh*)(Baker, 1987; Kornberg et al.,1985; Mohler & Vani, 1992), along the anterior posterior axis, just prior to the onset of gastrulation.

In embryos undergoing short or intermediate germ-band development, only some of the blastoderm cells coalesce to form a small patch of cells on the surface of the yolk mass, which represents the rudiments of only an anterior portion of the embryonic body plan. In these embryos, the remaining elements of the body plan form with the sequential addition of more posterior segments later, in concert with additional growth of the embryo within a posterior growth zone (Dearden & Akam, 2001; Lynch & Roth, 2011), the segment addition zone. Consequently, the volume of the egg corresponding to the embryo proper increases over the course of embryogenesis through the formation and growth of the initially absent more posterior segments. The distinction between short and intermediate germ-band development can be vague and, in fact, the development of the beetle *Tribolium castaneum* has been described as short germ-band in some publications (Sommer & Tautz, 1993; Nagy & Carroll, 1994) and as intermediate in others (Sulston & Anderson, 1996). For the purposes of this work, we define short germ-band-developing embryos as ones in which the initial primordium corresponds to the anterior portion of the head with all subsequent posterior segments being added sequentially, as in the grasshopper *Schistocerca gregaria* (Tear et al.,1990). By contrast, we define intermediate germ-band-developing insects as ones in which the initial primordium corresponds to the rudiments of the head and thoracic, or head, thoracic and anterior abdominal segments, having been specified simultaneously with the remaining posterior segments being added sequentially, as in the milkweed bug *Oncopeltus fasciatus* (Liu & Kaufman, 2004; Stahi & Chipman, 2016), the cricket *Gryllus bimaculatus* (Shinmayo et al., 2005; Donoughe & Extavour, 2016) and the beetle *Dermestes maculatus* (Patel et al., 1994; Xiang et al., 2015). It is important to note that in both short and intermediate germ-band-developing embryos, the period of segmental patterning extends beyond the onset of gastrulation, and that a number of posterior segments are added sequentially, with new segments being added “one by one” from a posterior segment addition zone.

To characterize the broad details of embryological development in myrmicine ants and determine the extent to which the major morphological events of embryogenesis are conserved between different ant species, we performed a partial analysis of embryonic development in *Atta texana*, *Mycoceparus smithii*, and *Solenopsis invicta*. Specifically, we compiled images representing a partial chronological series of successive developmental stages of embryogenesis ("developmental series") for these three ant species. For *A. texana*, we also characterized the expression of the two segmentation genes *en* and *wg,* owing to their conservation in the process of segmentation in insects and other arthropods (Patel et al., 1989; Nagy & Carroll, 1994; Brown et al., 1994A,B; Damen, 2002; Hughes & Kaufman, 2002). A previous study that included an analysis of early embryogenesis in the ants *Camponotus floridanus* and *Lasius niger* indicated that embryogenesis in these ants shared a feature of short germ-band development, namely that the embryonic primordium does not encompass the entirety of the anterior/posterior axis of the egg (Rafiqi et al., 2000). This was in contrast to the situation observed for two hymenopteran species in which embryogenesis has been examined, the honeybee (*Apis mellifera*) and the parasitic jewel wasp (*Nasonia vitripennis*). Both of these species exhibit a more typical long germ-band type of development, albeit modified in the case of *N. vitripennis* (Rosenberg et al., 2014).

Our findings indicate that the three myrmicine ants studied here do not exhibit the full complement of features characteristic of long germ-band development observed in other hymenopterans. Most notably, we also find that the embryonic primordium of each of these three ant species is quite small, with considerable growth of the embryo occurring prior to the establishment of segments, similar to what is seen in short and intermediate germ-band development. However, our gene expression study of *A. texana* does not support the addition of individual stripes of *en* and *wg* in concert with the growth of the embryo, as typically seen in short/intermediate germ-band development. Instead, following growth and elongation of the embryo, we observe a progressive addition of *en* and *wg* stripes of expression, from anterior to posterior, that serves to partition the embryonic germ-band into individual segments that occurs in the fully elongated embryo, more akin to what is observed in the long germ-band development of the mosquito *Anopheles stephensi* (Cheatle-Jarvela et al., 2021), a newly recognized mode of segmentation that has been termed progressive. Finally, we observe that in comparison to the other characterized hymenopterans *A. mellifera* and *N. vitripennis*, as well as *Drosophila*, the duration of ant embryogenesis is dramatically lengthened with much of this increase arising from the period of growth and the formation of segmentation within the fully elongated germ-band. In the case of *A. texana*, we find that the duration of embryogenesis of the progeny of founder queens is shorter, by approximately 25%, than the duration of embryogenesis in the progeny produced by queens in mature colonies. We discuss these findings in terms of the ant life histories and in particular with respect to selective pressures experienced by foundress queens during the establishment of their colonies.

## Materials and Methods

### Ant species

We studied embryo development in three ant species of the subfamily Myrmicinae (Hymenoptera, Formicidae), two fungus-growing ants (the leafcutter ant *A. texana* and the fungus-growing ant *M. smithii*), as well as the red imported fire ant *S. invicta*. Although these three species belong to the same subfamily (Myrmicinae, the largest subfamily of ants; Ward, 2007; Ward et al., 2015), they represent major differences in colony structure and ecology. For example, while both *A. texana* and *M. smithii* are fungus-growing ants within the same tribe (the Attini), *Atta* exhibits complex higher fungiculture, while *Mycocepurus* exhibits lower fungiculture (Schultz & Brady, 2008; Mueller, 2015; Mueller et al., 2017, 2018). Furthermore, *S. invicta*, the red fire ant, does not cultivate fungi. It is in the sister tribe Solenosidinii but is a highly invasive predatory ant that can cause serious damage to agriculture and natural environments (Ascunce et al., 2011). *S. invicta* also has a “social chromosome” that determines whether a colony is monogyne (single reproductive queen per colony) or polygyne (multiple reproductive queens per colony) in much the same way that the Y chromosome determines sex in mammals (Wang et al., 2013).

These three and species also exhibit differences in the extent of caste polymorphism. There are three different castes in *A. texana*: queen, male, and worker. Morphological castes in workers differ markedly in body size and head-to-body scaling (allometry) from small minims to large soldiers (Weber, 1972; Wetterer, 1999; Schultz & Brady, 2008). We collected *A. texana* foundresses after a mating flight on May 5, 2018 at Commons Ford Metropolitan Park in Austin, Texas, USA. Queens for the mature *A. texana* colonies used in this study were collected in 2014 after a mating flight in Fort Belknap, Texas, then colonies started from those queens were kept in our lab for several years. *Mycocepurus smithii*, is an unusual ant species that reproduces asexually by thelytokous parthenogenesis (Himler et al., 2009; Rabeling et al., 2009, 2011; Rabeling & Kronauer, 2013), in which diploid female offspring develop from unfertilized eggs (Fernández-Marín et al., 2005; Himler et al., 2009). There exist therefore only castes in a *M. smithii* colony, queens and monomorphic workers. Colonies of *M. smithii* were originally collected in 2001 from Gamboa, Panamá and were maintained under laboratory conditions as described previously (Rabeling et al., 2009; Kellner et al., 2013). Rabeling et al. (2011) documented that sexual populations of *M. smithii* appear to exist in the Amazonas region of Brazil, but not in the Panamanian populations where our lab colonies had been collected (e.g., no males have been found in our experimental clonal lineages kept in our lab since 2001).

While fire ant workers do not exhibit distinct morphological worker castes with respect to allometry, they do differ in total body size (Wheeler, 1991). Colonies of *S. invicta* exist in two forms, single-queen (monogyne) and multiple-queen (polygyne) colonies (Ross & Fletcher, 1985). Worker size is generally smaller in polygyne colonies than the workers in monogyne colonies (Greenberg et al., 1985). *S. invicta* colonies were collected in 2017 from Ecolab properties in Austin, Texas, USA. In this study, we only collected the embryos from polygyne colonies of the fire ants.

### Colony establishment and maintenance

*M. smithii* colonies were maintained as lab colonies using the methods of Sosa-Calvo et al. (2015). To establish mature lab colonies with large gardens, the wild-collected *A. texana* queens were allowed to found lab colonies using the methods of Marti et al. (2015). To compare embryogenesis in these mature colonies with embryogenesis of newly founded colonies, we also collected *A. texana* foundresses using the methods of Fang and Mueller (2019). Both fungus-growing ant species were given water *ad libitum* together with a mixture of polenta and finely ground oat flakes (ratio 3:1) as substrate for fungiculture. Attine colonies were kept in a temperature-controlled (but not humidity-controlled) laboratory room (22-24 °C; 12:12 [L:D]) in the Patterson Laboratories Building at the University of Texas at Austin. The colonies of *S. invicta* were maintained in a controlled environment (25 °C; 12:12 [L: D]; RH 60 ± 5%) in the Fire Ant Lab at Brackenridge Field Lab of the University of Texas at Austin. Each *S. invicta* colony was given *ad libitum* water, sugar water, and frozen crickets.

### Embryo collection

Because the egg-laying capacities of queens of the three species are different, we used different methods for egg collection. For *M. smithii*, because one queen can produce only 1-2 eggs in a 24-hour period (Fang et al. 2020), we pooled 10 queens and transferred them to the same Petri dish overnight for simultaneous egg collection. For *A. texana,* we placed a single queen into a round plastic container (5.5cm diameter; 3.7cm height) for egg collection, as described in detail by Marti et al. (2015) and Fang and Mueller (2019). A typical young foundress of *A. texana* lays 5-15 eggs in a 24-hour period, while a typical mature queen can lay 50-100 eggs within 24 hours. For both attine ant species, we added five workers and a small fragment of fungus (3 mm^3^) into the same container in which eggs were collected, to promote care of the eggs by the workers. Attine workers transfer freshly-laid eggs to the small fungal garden, which facilitates the detection and recovery of newly-laid eggs. We used a bottom-layer of 3% agar as the substrate in the containers to maintain humidity, but the smooth, clear agar bottom also allowed simple detection of any eggs. We added 15µl of diluted ampicillin (10 mg/ml) on a small piece of filter paper (about 2 x 2 cm^2^), which we placed on the agar plate during egg collection to minimize bacterial contamination of the fungus. For fire ants, we placed a single *S. invicta* queen, together with five workers, into a Petri dish with a bottom layer of 3% agar as the substrate, as described above. Because *S. invicta* queens oviposit at a high rate, we collected freshly-laid eggs 4 hours after introduction of the queen to the dish. For each of the three species investigated, clutches of eggs were collected daily (starting from the first 24 hrs, day 1), and the individual clutches of eggs were allowed to develop for a predetermined number of days of a developmental series. At collection, the eggs were fixed and stained as outlined below. Embryos obtained from these timed collections are shown in the figures displayed below.

### Embryo staging

Two other recent studies describe more detailed descriptions of the development from egg to adult in the pharaoh ant, *Monomorium pharaonis* (Rajakumar et al., 2023), and in the carpenter ant, *Camponotus floridanus* (Chen et al., in prep.), for which a partial analysis of embryogenesis has previously been reported (Rafiqi et al., 2020). For the sake of consistency, we have incorporated the staging system reported in Rajakumar et al. (2023) in the figures shown here.

### Embryo fixation

The protocol of embryo fixation was modified from Rafiqi et al. (2020). We moistened a fine brush with distilled water, then placed the embryos into a custom-made meshed basket. The basket was constructed by annealing a metal mesh to the cut bottom-end of a 50 ml conical centrifugation tube. To permeabilize the chorion of the eggs, we left the eggs on the mesh, which was then placed in a Petri dish. 10 ml of 15% bleach, diluted in water, were then poured into the Petri dish, followed by gentle agitation of the eggs by pipetting the solution up and down for 2 minutes to wash the eggs. After permeabilization, we rinsed the eggs for 30 sec in distilled water to eliminate any residual bleach. The eggs were then treated with 0.3% Triton X-100 for 5 min on ice, then transferred in their baskets to boiling 0.3% Triton X-100 for 20 sec. Eggs were then rinsed quickly in ice-cold 1x phosphate buffered saline (PBS), then transferred into ice-cold 0.1% Tween-20 diluted with PBS (PBST) for 5 min. For more convenient manipulation, we then transferred the eggs to a 1.5 ml Eppendorf tube and rinsed them three times in 0.1% PBST. Following the third rinse, the eggs were incubated in 0.1% PBST on ice for 1 hour. To permeabilize vitelline and embryonic membranes, the eggs were treated with Proteinase K (20 µg/ml stock solution in ethanol, diluted 500x in 0.1% PBST) for 5 min, followed by immersion in 1x glycine (20 mg/ml stock solution in EB buffer, diluted 10x in 0.1% PBST) for 5 min on ice. The eggs were then rinsed three times with PEM buffer (recipe for 250mL PEM buffer is 8.66g PIPES, 0.19g EGTA, and 0.03g MgSO_4_ in distilled water). The eggs were then moved into a new 1.5 ml Eppendorf tube with the fixation solution (400µl PEM, 60µl 37% formaldehyde, and 500µl Heptane), then the samples were rotated (180 rpm) at room temperature for 25 min. After removing the post-fix solution, the eggs were incubated briefly in 200µl distilled water for one minute. Finally, we washed the eggs three times in ice-cold methanol (MeOH). Fixed embryos were stored in MeOH at -20°C.

### DAPI staining

To rehydrate the fixed samples, we treated the eggs at room temperature with 75%, 50%, and 25% MeOH (diluted with 0.1% PBST) for 5 minutes each. The eggs were then rinsed three times with 0.1% PBST, then washed for 10 min with 0.1% PBST with shaking (100 rpm) at room temperature. To ensure quality staining, we applied a second permeabilization to the eggs, as follows: proteinase K treatment for 5 min was followed by 1x glycine for 5 min, followed by rinses with 0.1% PBST. Post-fixation (in 135µl formaldehyde with 865 µl 0.1% PBST) was performed for 25 min, followed by another 5 min wash in 1x glycine. To remove the post-fixation solution, we rinsed the sample again three times for 5 min in 0.1% PBST. The eggs were then immersed in a DAPI solution (1uL DAPI with 999µl 0.1% PBST) for 30 min, then washed with 0.1% PBST for 5 min. Eggs were mounted in 75% glycerol (diluted with 0.1% PBST), followed by incubation in a series of solutions of increasing proportions of glycerol diluted in 0.1% PBST as follows: 25% glycerol for 5 min; 50% glycerol for 30 min; 75% glycerol overnight at 4 °C.

### Whole-mount immunohistochemistry

To characterize the expression of the Engrailed (En) protein during segmentation, we treated the dissected embryos with monoclonal anti-engrailed (clone 4D9 from the Developmental Studies Hybridoma Bank, DSHB) as the primary antibody and goat anti-mouse (Alexa Fluor 488) as the secondary antibody. The entire antibody staining procedure takes 2 days. On Day 1, the dissected embryos were rehydrated and subjected to post-fixation prior to incubation with the monoclonal primary antibody on Day 1. The embryos were then blocked overnight in 5% normal goat serum (NGS) (diluted with 0.1% PBST), then treated with 4D9 at 4 °C. Concentration for 4D9 was 5µg/µL (diluted 4D9 in 1% NGS; 1% NGS was diluted with 0.1% PBST). On Day 2, embryos were washed with 1% NGS (diluted with 0.1% PBST) 4 times at room temperature, then treated with the secondary antibody (diluted 1:750 in 1% NGS) for 2 hours at room temperature. This was followed by washing of embryos with PBST four times, for 15 min each. The embryos were then mounted for imaging as described above for DAPI staining.

### Hybridization chain reaction RNA fluorescence in situ hybridization (HCR RNA-FISH)

To characterize the expression of *wingless* mRNA throughout embryonic development, we performed the HCR RNA-FISH method (Choi et al., 2018) on whole mount preparations of embryos. To visualize *Atta wingless* mRNA, 20 probes were generated against *A. colombica wingless* (XM_018204929) utilizing the Molecular Instruments probe design services. HCR was performed using the manufacturers buffers and protocol (Choi et al., 2018).

### Microscopy

Fluorescent microscopy images were obtained on a Zeiss Axiovert Fluorescent Light Microscope at the Microscopy and Imaging Facility at the University of Texas at Austin (http://sites.cns.utexas.edu/cbrs/microscopy). Images of embryos that had been subjected to *in situ* hybridization were taken using a Leica MZ16 Stereomicroscope fitted with a DFC420 digital camera; images of immunohistochemistry were taken using a Nikon confocal microscope C2/C2si fitted with Ti2-LAPP Ti2 laser application system. Images of embryos obtained at McGill University were taken on a Zeiss Discovery V12 stereomicroscope with Zeiss Axiovision software.

## Results

### Embryogenesis of three myrmicine ants

To test whether the major morphological events of embryogenesis are conserved between different ant species, we performed a partial analysis of embryonic development in *Atta texana*, *Mycoceparus smithii*, and *Solenopsis invicta*. In *A. texana*, embryonic development of the progeny of foundress *A. texana* queens lasts about 15 days at 25 ± 1 °C (Fig. 1). However, as will be described below, embryogenesis in the progeny of mature queens lasts for 20 days. In the progeny of founder queens, the zygote nucleus has undergone sufficient mitotic divisions by the end of Day 2 to generate a cellular blastoderm embryo (stage 5) (Fig. 1b). On Day 3, later in stage 5, many of the nuclei are visualized at a posterior ventral position in the egg in the embryonic primordium (Fig. 1c). On Day 4 (stage 6), folds are observed in the developing embryo suggesting that the onset of gastrulation is occurring (Fig. 1d). This can best be seen viewing embryos from the ventral side. Following this stage, the embryo enters a period of pre-segmentation growth and extension (elongation) along the anterior-posterior axis, which lasts from Day 4 to Day 7 (stages 6 through stage 8)(Fig. 1d-g). At the end of the pre-segmental elongation phase, the future head lobe bulges out on Day 8 (stage 8)(Fig. 1h), which is followed by morphological signs of progressive segmentation appearing along the anterior-posterior axis. On Day 9, the three gnathal segments are observed, labeled mandibular (md), maxillary (mx), and labial (lb) in Fig. 1i. On Day 10 (Fig. 1j), the future thoracic region starts to form, and the three distinguishable future thoracic segments can be observed on Day 11 (Fig. 1k). After development of the thoracic segments, the morphologically distinct segments are visible throughout the abdomen on Day 12 (Fig. 1l). On Days 13 and 14, germ-band retraction proceeds gradually (Fig. 1m-n). On Day 15, the 1st-instar larva hatches (at 25 ± 1 °C) (Fig. 1o) to complete embryogenesis of eggs laid by a foundress queen in a period of 15 days. In contrast, complete embryogenesis of eggs laid by a mature queen takes 20 days (see details below).

**Figure 1.**
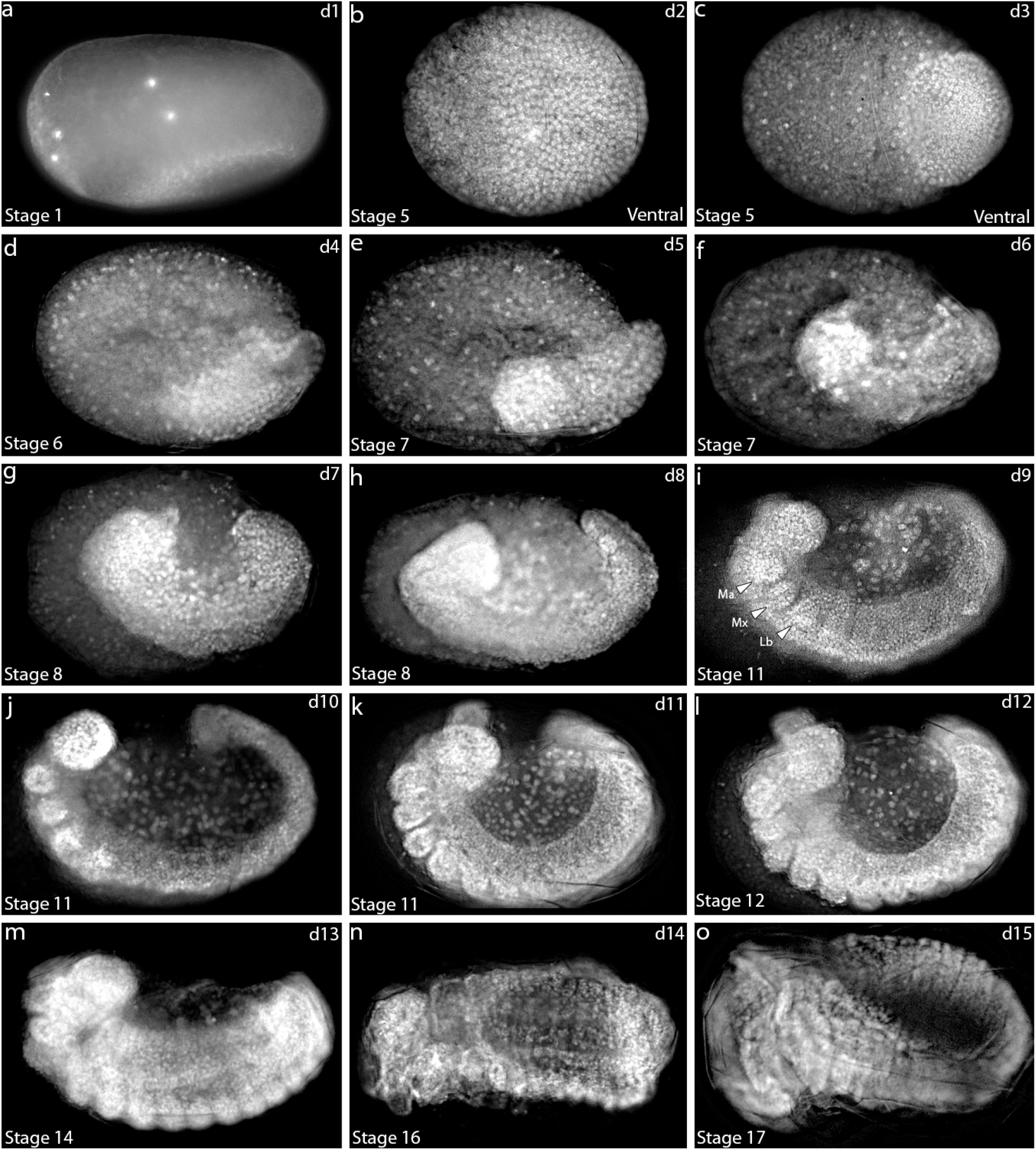
A series of images (“developmental series”) of embryos produced by *Atta texana* foundress queens, covering day 1 to day 15 after egg deposition (the egg stages). The embryos are staged (from 1 to 17) day (d)1 to day 15 (d15) of development as described in Pontieri et al. (2020). The panels show a series of images corresponding to successive developmental stages of embryos. Fixed embryos that have been removed from their eggshells are stained with the nuclear stain DAPI. The blastoderm stage occurs 24-48 hours after oviposition. A condensed embryonic primordium appears posteriorly on day 3. The germ-band undergoes pre-segmental elongation on days 4 (d) through day 7(g), and the first segment (future head lobe) appears on day 8(h). A clear separation of the future head and gnathal segments develops on day 9; the three gnathal segments developing on day 9 are the mandibular (md), maxillary (mx), and labial (lb) segments (see arrowheads, panel i). The process of embryogenesis for the progeny of foundress females lasts 15 days at 25 ± 1 °C, ending with hatching of the 1st-instar larva on day 15 (o); In contrast, the egg stage for the progeny of mature queens is slower and lasts 20 days (see Fig. 7).

In *M. smithii*, embryogenesis lasts 19-20 days at 25 ± 1 °C (Fig. 2). The first mitotic division occurs within 24 hours (Fig. 2a) and the blastoderm stage begins on Day 2 (Fig. 2b). On Day 3-4, nuclei are visualized at a posterior ventral position in the embryonic primordium (Fig. 2c-d). Between Days 5-6, distinctive folds develop (Fig. 2e-f), which are best visualized in the ventral view (Fig. 2e), and which likely show that the embryo is undergoing gastrulation. Germ-band growth and pre-segmental elongation occur between Day 6 to Day 10 (Fig. 2f-j). The first body segment (i.e., future head lobe) is observed on Day 9, and the germ-band starts to curve from the lateral side and extend towards the posterior side (Fig. 2i). On Day 10, bulges corresponding to the three gnathal segments begin to form at the anterior part of the germ-band (Fig. 2j). Between Days 11-15, the future thoracic and abdominal segments appear progressively (Fig. 2k-o). Germ-band condensation (retraction) occurs between Days 16-19. Between Days 19-20, the 1st-instar larva hatches (at 25 ± 1 °C) (Fig. 2p-s).

**Figure 2.**
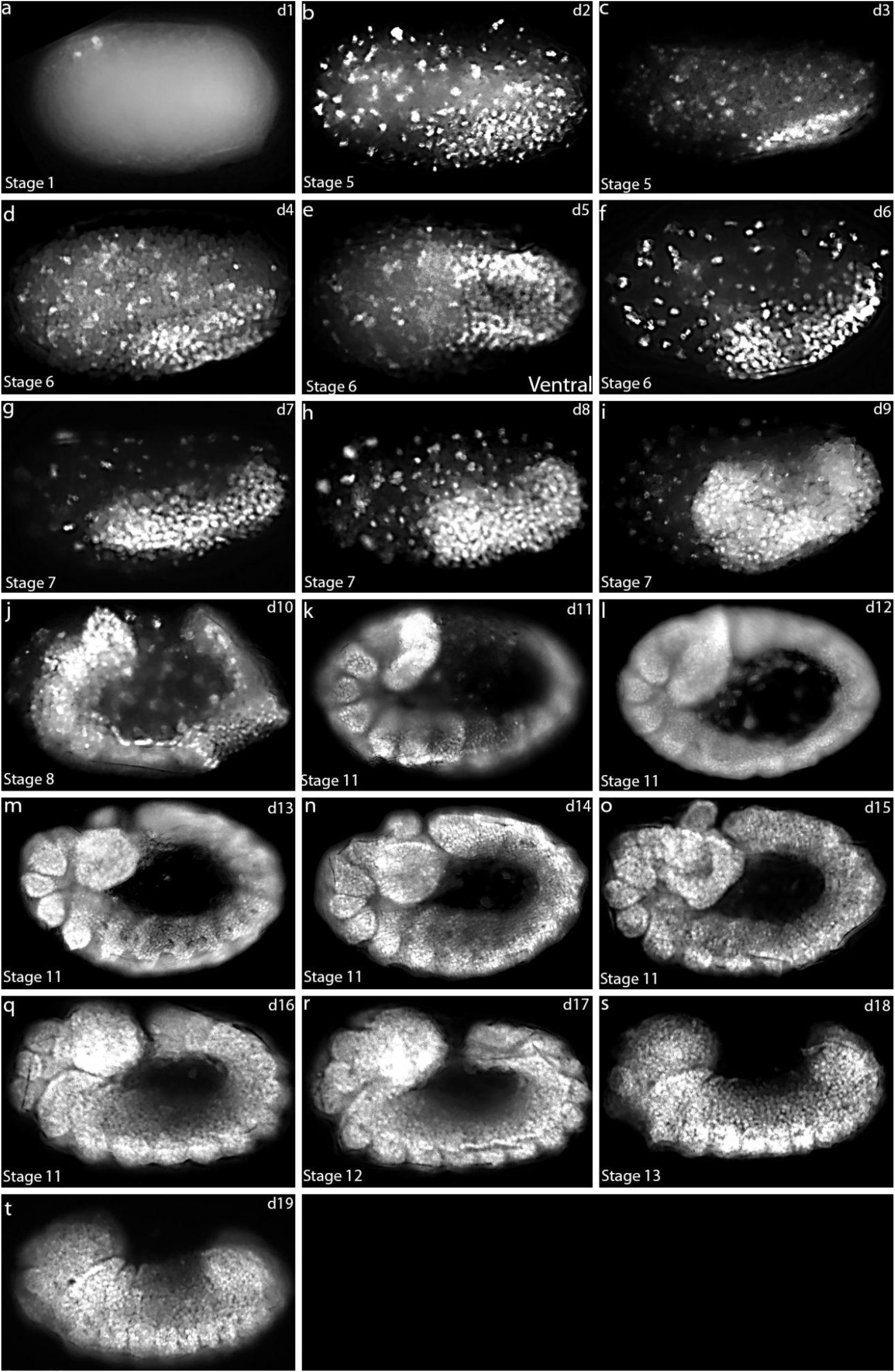
Developmental series of *Mycocepurus smithii* embryos, covering day 1 through day 19 following egg deposition (Stage 1 through Stage 14). Fixed embryos that have been removed from their eggshells and stained with DAPI are shown. The blastoderm stage occurs 24-48 hours after oviposition (panel b). A heart-shaped germ-band develops on day 5(e). The germ-band undergoes pre-segmental elongation on days 6-10 (f-j), and the first segment (future head lobe) begins to appear on day 9 (i). A separation of the future head and the three gnathal segments begins to appear on day 10 (j). Thoracic and abdominal segments form sequentially and increase in number from day 11 to day 15. Embryonic segmental condensation occurs from day 18 to day 19 (s,t). The total egg stage lasts 19-20 days at 25 ± 1 °C, ending with hatching of the 1st-instar larva on day 19-20.

Embryos of *A. texana* and *M. smithii* exhibit comparable periods of development at 25 ± 1°C: 15 days (eggs laid by foundress queen) or 19/20 days (eggs laid by mature queen) in the case of *A. texana* versus 19-20 days for *M. smithii*. The similar rates of development permitted relatively direct comparisons of developmental landmarks between these two species. In both species, multiple nuclei emerge on the surface of the embryos during the blastoderm stage (Fig. 3a, e). As development continues, the embryonic primodium takes on a distinctive heart-like shape (Fig. 3b, f). During embryonic gastrulation and elongation, the germ-band begins to curve and extend towards the posterior side (Fig. 3 c, g). Finally, the morphologically distinct future body segments form during the segmentation phase (Fig. 3d, h).

**Figure 3.**
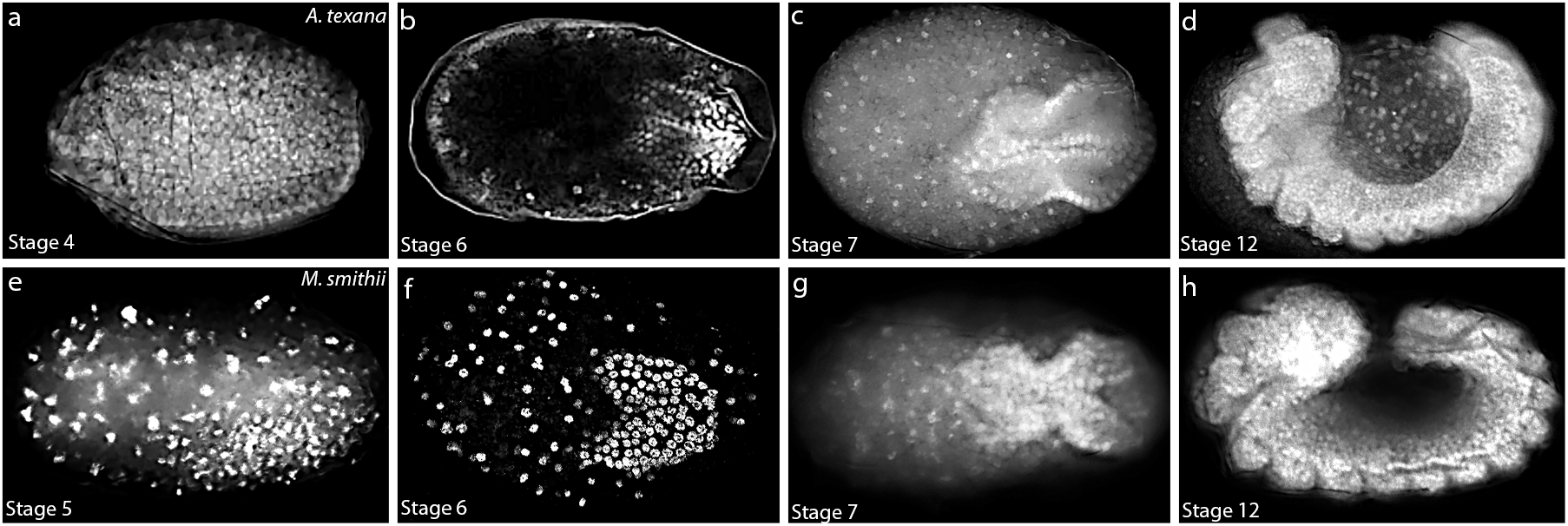
Comparison of embryogenesis in *Atta texana* (a-d) and *Mycocepurus smithii* (e-h). Multiple nuclei emerge on the surface of the embryos during the blastoderm stage (a, e). The heart-shaped germ-band forms towards the posterior end of the embryos (b, f). The germ-band begins to curve laterally and extend towards the posterior end during embryonic gastrulation and elongation (c, g). Morphologically distinct segments form subsequently (d, h).

Embryos of *Solenopsis invicta*, a species that belongs to the tribe Solenopsidini (sister clade to the tribe Attini that includes the fungus-growing ants; Ward et al., 2015), complete embryogenesis in a considerably shorter time than those of those of *A. texana* and *M. smithii*. The egg stage of *S. invicta* lasts 9-10 days at 25 ± 1 °C (Fig. 4) with the blastoderm appearing on Day 2 (Fig. 4b). Because of the more rapid development of *S. invicta* embryos, we were not able to identify or visualize embryos at the heart stage or early in gastrulation. Between Days 2-3 of *S. invicta* embryogenesis, the germ-band forms and extends rapidly (Fig. 4d,e). The future head lobe forms on Day 4 (Fig. 4f), and the remainder of the future body segments develop on Day 5 (Fig. 4g,h). The process of embryonic condensation extends from Day 6 to Day 8 (Fig. 4i,j). On Days 9 or 10, the 1st-instar larva hatches (at 25 ± 1 °C) (Fig. 4k-l). Together, our observations show that the major developmental stages were observed to be conserved in the three species, though accelerated in *S. invicta*.

**Figure 4.**
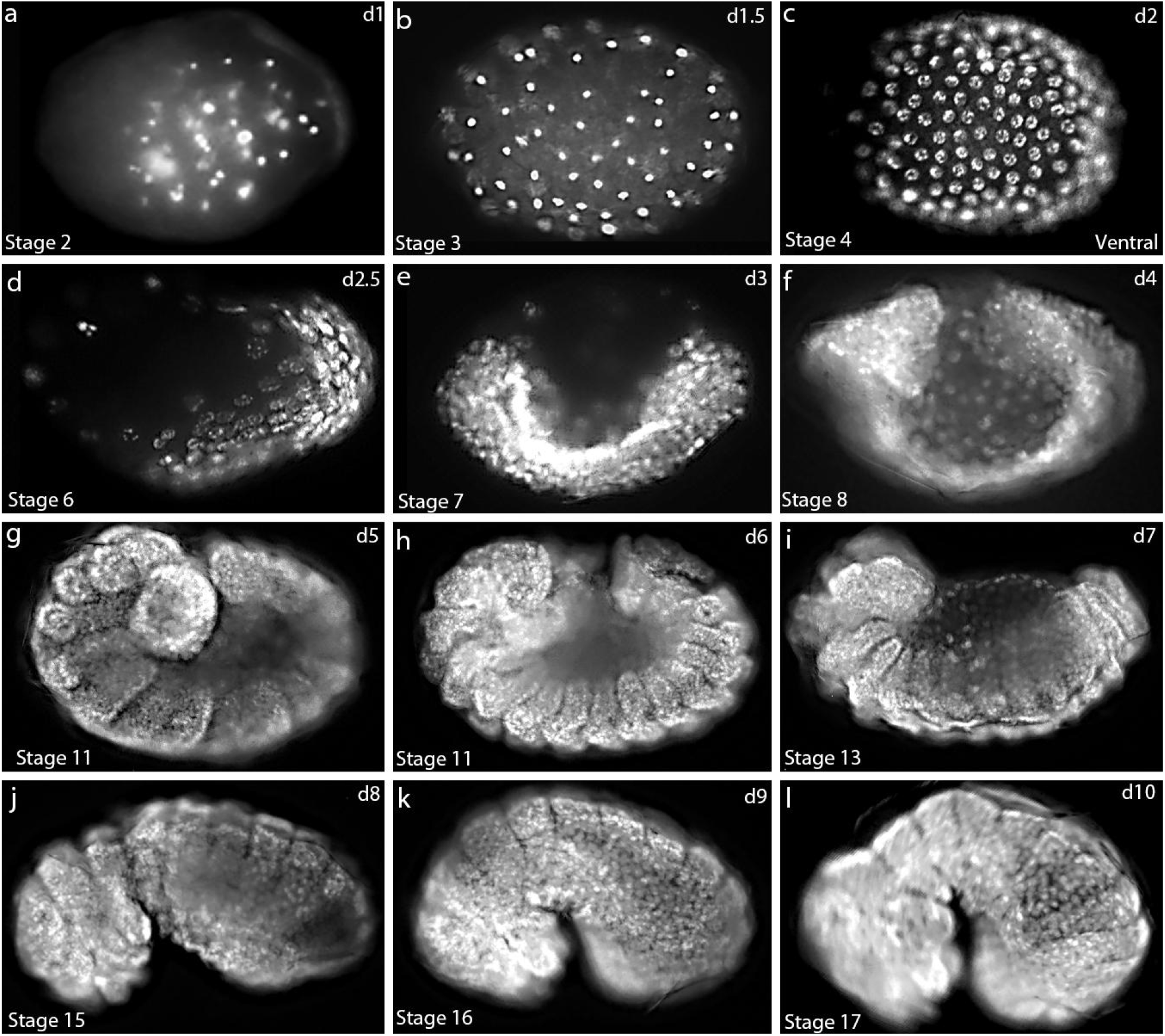
Developmental series of *Solenopsis invicta* embryos, from day 1 to day 10 following egg deposition (Stage 1 to 17). Fixed embryos that have been removed from their egg shells are stained with DAPI. The blastoderm stage occurs 24-48 hours after oviposition (panels a-c). The germ-band undergoes pre-segmental elongation over days 2-4 (d-f). The first segment (future head lobe) appears on day 4 (f), and segmentation occurs during days 5-6. The germ-band condensation occurs on days 7 and 8 (i,j) with hatching on day 9 or 10. The total egg stage lasts 9-10 days at 25 ± 1 °C.

### Expression of the segmentation gene products *wingless* mRNA and Engrailed protein

In *Drosophila*, formation of separate segments along the anterior-posterior axis of the embryo is controlled, in part, by Engrailed and the Wingless signal transduction pathway, both of which act to define the boundaries between segments and the polarity of cells contained within the individual segments that comprise the anterior-posterior axis (DiNardo et al., 1988, 1994; Larsen et al., 2008; Swarup & Verheyen, 2012). More accurately, these “segment polarity” genes define the boundaries between and the polarity of cells within parasegments (Fjose et al., 1985; Larsen et al., 2008; Swarup & Verheyen, 2012). In *Drosophila* embryos, the *wingless* (*wg*) gene is expressed in the most posterior cell of a forming parasegment, and the *engrailed* (*en*) gene is expressed in the anterior cell of the adjacent posterior forming parasegment (DiNardo et al., 1994; Larsen et al., 2008; Swarup & Verheyen, 2012). The roles of *en* and *wg* in the formation of segments is conserved in short germ-band-developing insects as well as the other arthropod groups (Farzana & Brown, 2008; Simonnet et al., 2004; Damen, 2007).

In long germ-band-developing *Drosophila*, fourteen stripes of *en* and *wg* expression can be detected during gastrulation. These stripes first appear over a period of about an hour and a half, from the cellular blastoderm stage to the onset of gastrulation, at discrete positions along the anterior-posterior axis of the germ-band, with engrailed stripe 2 forming first, with even-numbered stripes appearing before odd-numbered stripes (Weir & Kornberg, 1985; DiNardo et al., 1988) and with a general anterior-to-posterior progression. Engrailed is widely conserved among arthropods and a single monoclonal antibody, MAb 4D9 (Patel et al., 1989), has often been used to examine the pattern of segmentation in different insect and arthropod species. In contrast to the situation in *Drosophila*, in short germ-band-developing insects, (e.g. the grasshopper *Schistocerca gregaria*) the stripes of Engrailed protein appear one at a time, starting at the onset of gastrulation. The remaining Engrailed stripes form within a posterior growth zone and are added one by one as the embryo adds new segments via growth (Dearden & Akam, 2001). In the beetle *Tribolium castaneum*, 5 Engrailed stripes are present at the beginning of germ-band elongation and newly-forming Engrailed stripes anticipate the formation of each new segment, as they appear sequentially from the anterior to the posterior of the elongating germ-band (Brown et al., 1994A, B).

Owing to the conservation of the involvement of segment polarity genes in the formation of segments in insects and other arthropods, we elected to investigate the dynamics of Engrailed and *wingless* expression in embryos produced by mature *A. texana* queens in order to better understand this process and to define the germ-band classification into which this ant species fall. In *A. texana*, prior to localization of the embryonic primordium at the posterior end of the egg, no *wg* mRNA is detected (Fig. 5a-c). (Fig. 5d). Expression of *wg* first appears at stage 6 during which the signal can be detected at both the anterior and posterior ends of the embryonic primordium (Fig. 5d). As the embryo begins to elongate, additional foci of *wg* mRNA can be detected along the anterior-posterior axis of the embryo (Fig. 5e-f). The appearance of *wg* expression in locations corresponding to the primordia of the mandibular, maxillary, and labial, and three thoracic segments is first observed at the beginning of stage 8 (Fig. 5g). As stage 8 continues, the gnathal (Ma, Mx, and Lb) and thoracic (T) stripes become more conspicuous and additional stripes of *wg* mRNA corresponding to abdominal (A) segments appear and increase in number progressively (Fig. 5h-l) until the complete complement of ten abdominal stripes are discernable (Fig. 5l). The *wg* bands become more distinctive in stages 9 and 10 (Fig. 5m-p), followed by the formation of morphologically distinct abdominal segments (Fig. 5q-s). From stages 12 -16 (Fig. 5t-x), *wg* expression was detected in each body segment during the process of germ-band retraction (condensation). We also examined Engrailed (En) protein expression in eggs laid by a mature *A. texana* queen from Day 9.5 to Day 11 after oviposition (Fig. 6). Consistent with *wg* expression (Fig. 5), following growth and germ-band extension, 6, 8, and 13 forming (Fig. 6a) and distinct stripes (Fig. 6b,c) of Engrailed protein could be seen on Day 9.5, Day 10, and Day 11, respectively.

**Figure 5.**
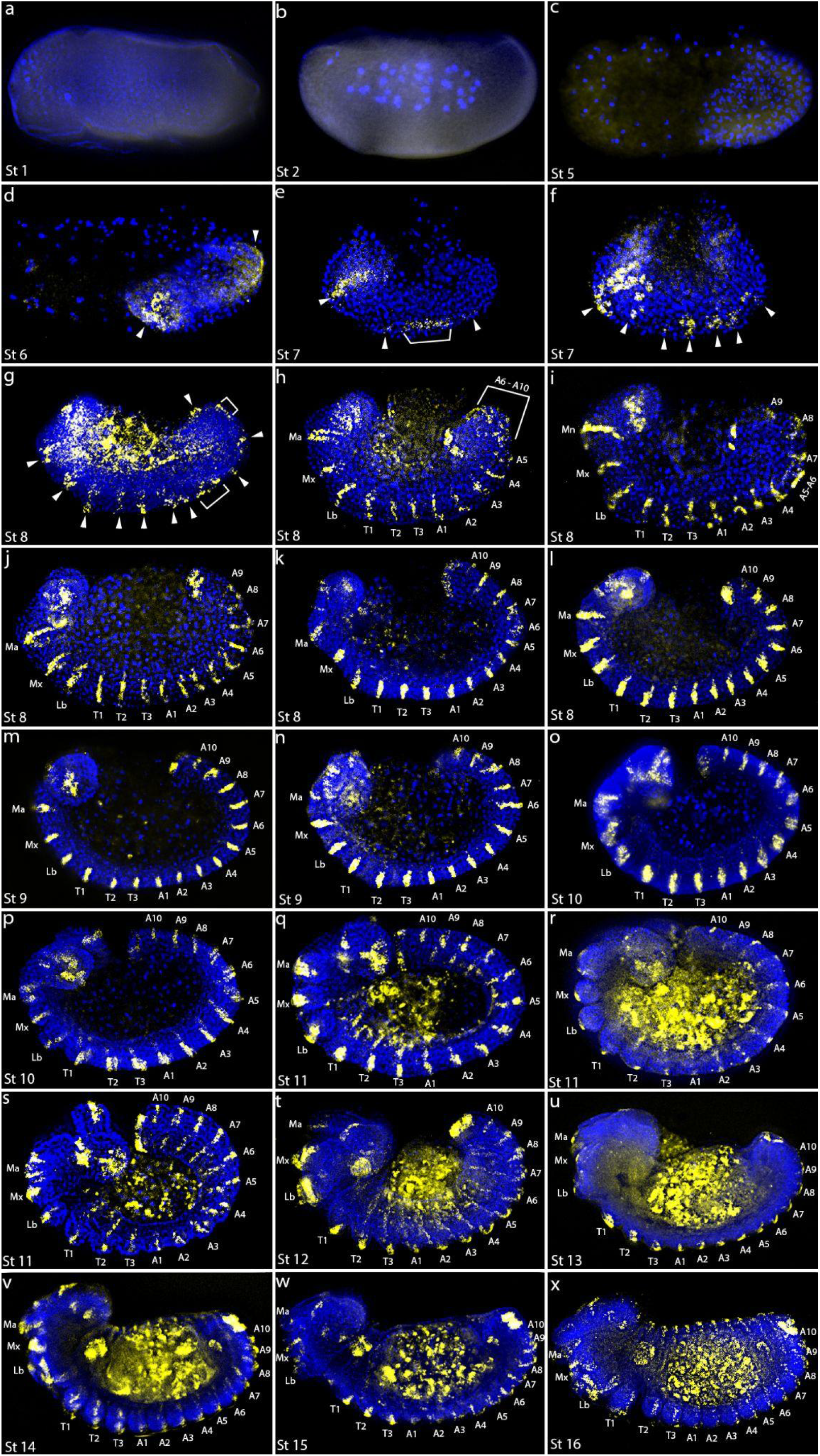
Developmental series of embryos expressing *wingless* (*wg*). (a-x) *A*. *texana* embryos stained with *wingless* (*wg*) HCR probe (yellow) counterstained with the nuclear stain DAPI. (a) Stage 1 syncytial stage lacking *wg* staining. (b) Stage 2 syncytial stage lacking *wg* staining. (c) Stage 5 cellular blastoderm stage lacking *wg* staining. (d) Stage 6 cellular blastoderm stage. (e,f) Stage 7 gastrulation stage. (g-l) Stage 8 germ-band extension stage. (m,n) Stage 9 germ-band extension stage. (o, p) Stage 10 germ-band extension stage. (q-s) Stage 11 segmentation stage. (t) Stage 12 germ-band retraction stage. (u) Stage 13 germ-band retraction stage. (v) Stage 14 dorsal closure stage. (w) Stage 15 dorsal closure stage. (x) Stage 16, embryo straightens. White arrows (d-g) indicate early *wg* stripes. White brackets (e, g, h) indicate grouped unresolved *wg* stripes. Abbreviations and the structures that they designate are as follows: Ma, mandible; Mx, maxilla; La, Labium; T1, 1^st^ thoracic segment; T2, 2^nd^ thoracic segment; T3, 3^rd^ thoracic segment; A1, 1^st^ abdominal segment; A2, 2^nd^ abdominal segment; A3, 3^rd^ abdominal segment; A4, 4^th^ abdominal segment; A5, 5^th^ abdominal segment; A6, 6^th^ abdominal segment; A7, 7^th^ abdominal segment; A8, 8^th^ abdominal segment; A9, 9^th^ abdominal segment; A10, 10^th^ abdominal segment. In all panels the embryos are oriented with their anterior end to the left, posterior end to the right, dorsal side up, and ventral side down.

**Figure 6.**
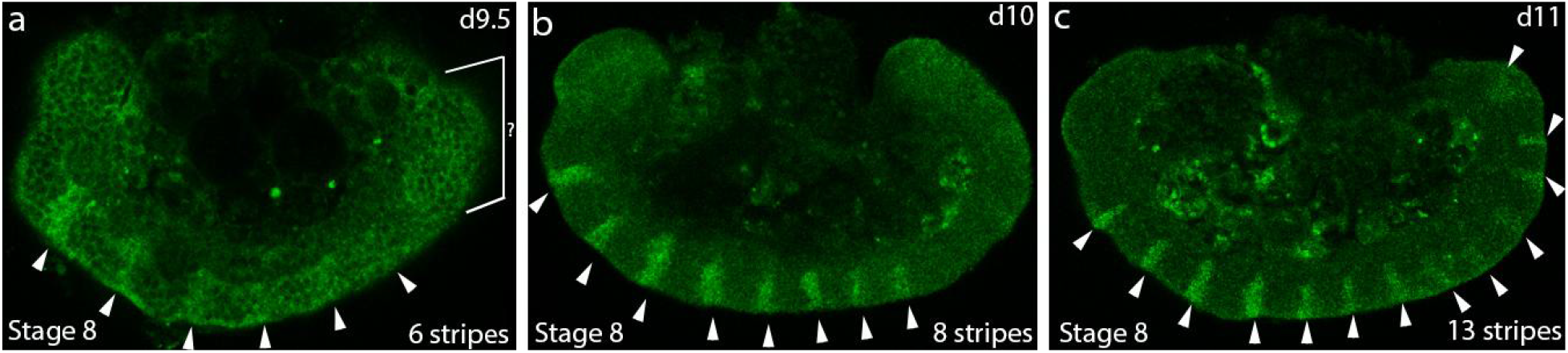
Engrailed (En) protein expression in embryos of *Atta texana* on day 9.5 (a), day 10 (b), and day 11 (c) after egg deposition. Eggs were collected from a mature *Atta texana* queen and maintained at 25 ± 1 °C. The embryos are oriented to show the anterior on the left. The number of stripes of En expression increases with time. On day 9.5 (a), 6 forming bands enriched in En protein can be discerned (see arrowheads); on day 10 (b) 8 condensed bands appeared; on day 11, 13 stripes were discernible.

### Duration of embryogenesis in the progeny of *Atta texana* foundress versus mature queens

Following the nuptial flight, newly fertilized queen ants typically excavate a tunnel leading to a small chamber, in which she seals herself and lays her first batch of eggs which she will need to rear without the benefit of workers (Hölldoper and Wilson, 1990). In ant species engaging in fungiculture, founder queens are additionally tasked with the responsibility of maintaining the fungal culture that will ultimately provide sustenance to the colony. In most cases, newly mated queens do not leave their nests to forage, a behavior that reduces her risk of predation, desiccation, or other injury. This requires that the founder queen rears her first batch of worker offspring using stored body resources (Martinez & Wheeler, 1994), which become depleted over the period during which she rears her first brood (Wheeler & Buck, 1996). In addition to these physiological adaptations, the founder queens exhibit brood care behavior that they will abandon when her first brood reaches adulthood (Cassill, 2002). These changes in ant queen physiology and behavior led us to ask whether then embryos of the first brood of *A. texana* foundress queens exhibit developmental acceleration to cope with the time constraints of founding an ant colony. We therefore timed the duration of development of embryos by foundress queens and mature queens as outlined in the Materials and Methods section. When maintained at the same temperature (25 ± 1 °C), embryos produced by foundress queens developed 25% more rapidly than the embryos produced by mature queens (Fig. 7). For the foundress-laid eggs and mature queen-laid eggs, the duration of embryogenesis from oviposition to hatching first instar larva was 15 and 20 days, respectively. To further characterize developmental trajectories between the two types of embryos, we examined the blastoderm, heart-shaped, pre-segmental elongation, and segmentation stages, which represent four embryonic developmental landmarks that we have used in our analyses of ant embryogenesis (Fig. 7). There were no apparent differences in the durations of the blastoderm (Day 2) and heart-shaped stages (Day 3) exhibited by embryos from the two types of queens. However, foundress-laid eggs exhibited shorter durations of the pre-segmental elongation and the segmentation stages. Embryonic pre-segmental elongation started on Day 4 in the embryos produced by both foundresses and mature queens. However, the duration of this stage was four days in the progeny of founder queens and seven days in the embryos from mature queens, such that segmentation began on Day 8 in eggs laid by foundress queens, but did not start until Days 10-11 in eggs laid by mature queens (Fig. 7).

**Figure 7.**
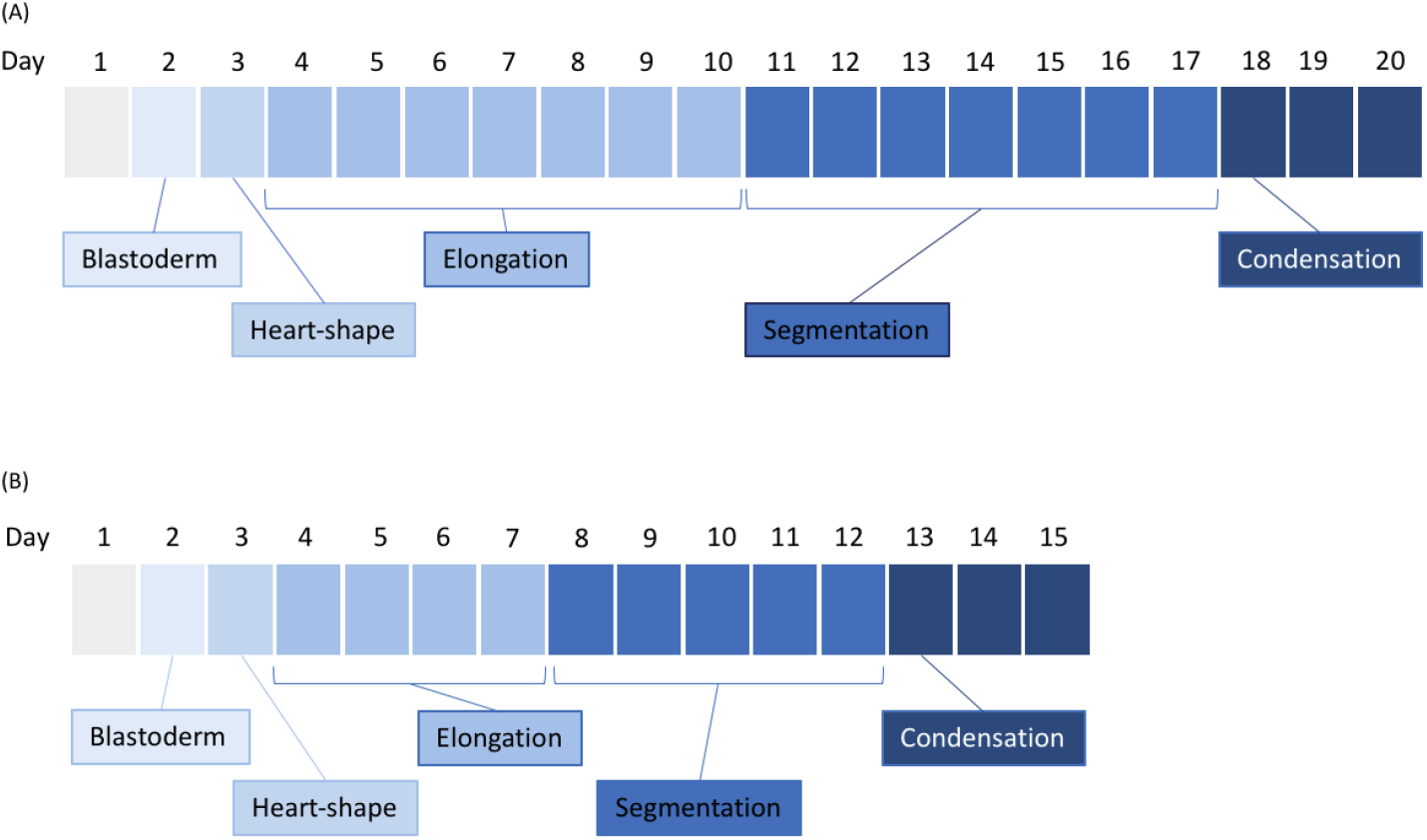
Developmental trajectories of (A) mature queen-laid embryos and (B) foundress-laid embryos of *Atta texana* at 25 ± 1 °C. Embryos in eggs laid by a foundress queen develop 25% faster than embryos from a mature queen, requiring 15 days versus 20 days for development from oviposition to hatching of the 1^st^-instar larvae. Blastoderm development occurs on day 2 in embryos from both sources, but subsequent development is more rapid in embryos produced by a foundress queen. Among the four embryonic developmental stages, the differences in duration of embryogenesis primarily affect the stages of pre-segmental elongation (denoted as “elongation”) (4 days versus 7 days post-egg deposition, respectively, by foundress versus mature queens) and segmentation (5 days versus 7 days post-egg deposition).

Comparing the two types of embryos on Day 9, a foundress-produced embryo showed a well-developed future head lobe, as well as mandibular, maxillary, and labial (lb) gnathal segments from anterior to posterior (arrowheads in Fig. 8a). In contrast, for embryos produced by mature queens, on Day 9 the future head was beginning to form, while the three gnathal segments had not yet started to develop (Fig. 8c). On Day 12, foundress-derived embryos showed the three gnathal segments as well as three well-defined segmental anlage of the future thorax (from anterior to posterior, arrowheads in Fig. 8b), but there was no obvious morphological thoracic segmentation by that time in the embryos derived from mature queens (Fig. 8d). In fact, these embryos had not developed as far as embryos from founder queens had by day 9.

**Figure 8.**
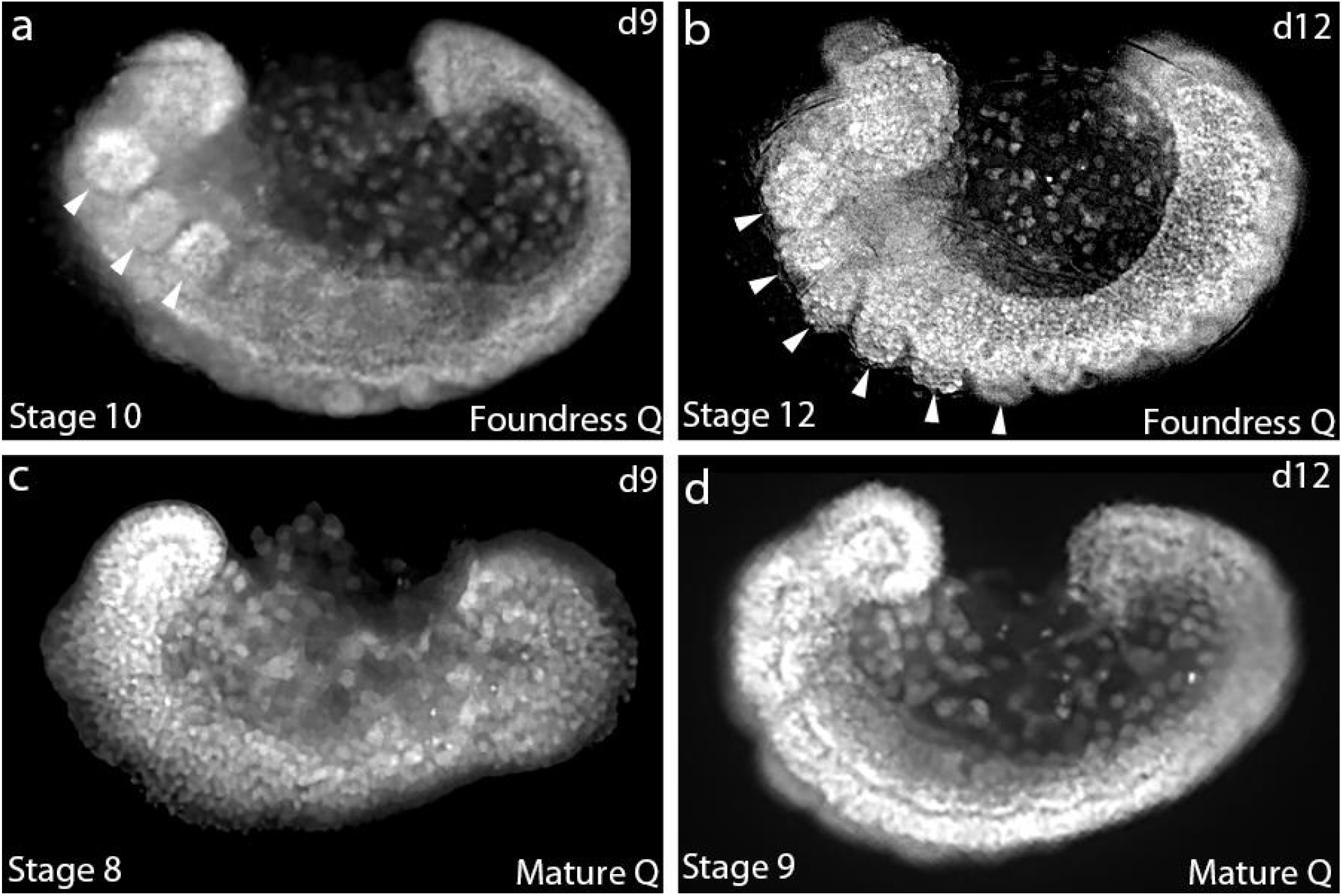
Comparative morphology of 9-day-old embryos produced by an *Atta texana* foundress queen (a) and a mature queen (c), and 12-day-old embryos produced by an *Atta texana* foundress queen (b, panel l from Fig. 1) and a mature queen (d). The eggs are oriented to show the anterior end to the left. In an egg of a more rapidly-developing worker laid by a foundress queen (a), the future head lobe and gnathal segments (three arrowheads in a) are already formed by day 9. In contrast, an embryo produced by a mature queen on day 9 exhibits only partial development of the future head lobe, and no morphologically distinguishable thoracic segmentation (c). (b) Formation of the future head lobe and morphologically-distinguishable body segments by a 12-day-old embryo from a foundress queen (with the 6 arrowheads corresponding to the gnathal and three thoracic segments) compared to (d), a 12-day-old embryo from a mature queen displaying a well-developed future head lobe but lacking obvious morphologically-distinguishable body segments.

## Discussion

Our study represents an initial characterization of segmentation of three ant species in the subfamily Myrmicinae, two fungus-gardening ants (the leaf-cutter ant *A. texana* and a so-called primitive fungus-growing ant *M. smithii*), and the red imported fire ant *S. invicta*. One goal was to use embryological characteristics and the expression of the segment polarity genes Engrailed and Wingless to categorize in these species the mode of embryogenesis, short, intermediate, or long germ-band (Davis and Patel 2002). Interestingly, the ant embryos examined do not exhibit the conventional constellation of features typical for long germ-band development as observed in the hymenopteran species *Apis mellifera* and *Nasonia vitripennis* (Lynch et al., 2012; Wilson et al., 2014; Cridge et al., 2017). Most conspicuously, the apparent embryonic primordia formed by these ant embryos comprise only a small proportion of the egg volume, and considerable growth of the embryo appears to occur over the course of development such that the embryo ultimately fills the entire volume of the egg. These are embryological features that are typically observed in insects undergoing short/intermediate germ-band types of development (Dearden & Akam, 2001; Handel et al., 2005). Visually, this early partitioning of embryonic and extraembryonic cells in the ants appears to be quite distinct from what is occurring in the typical long germ-developing insect such as *D. melanogaster*.

To determine when the germ-band generates its full complement of segments, we examined expression of the segmentation genes *engrailed* and *wingless* (Cabrera et al., 1987; Dearden & Akam, 2001; Swarup & Verheyen, 2012; Vellutini & Hejnol, 2016). In *Drosophila*, the canonical long germ-band insect, a full complement of stripes of *engrailed*/*wingless* expression forms prior to the completion of gastrulation (Baker, 1988; Weir & Kornberg, 1985). Moreover, fate mapping studies of *Drosophila* indicate that a full complement of segments has been defined molecularly in the cellular blastoderm stage embryo, prior to the onset of gastrulation (Akam, 1994; Lohs-Schardin et al., 1979; Patel et al., 1992). The strict association between long germ-band development and the nearly simultaneous segmentation prior to gastrulation is exhibited in its most extreme case in *Drosophila melanogaster* and other brachyceran (“true”) flies. Studies of additional holometabolous insects demonstrate that this association is not strict. Although the closely-related Nematoceran flies (mosquitoes, gnats, and midges) would be classified as long germ-band-developing insects based on the proportion of blastoderm cells that comprise the embryonic primordium, these embryos delay the specification of posterior segments (Cheatle-Jarvela et al., 2021). However, in contrast to sequential segmentation, the addition of new segments from anterior to posterior, does not depend upon the formation of segments from within a segment addition zone. Rather, this mode of segment addition, which has only been described in long germ-band insects, results from the progressive appearance of pair rule and segment polarity expression stripes along the anterior-posterior axis of the germ band. This mode of segmentation has been termed progressive segmentation and represents an intermediate mode between the sequential and simultaneous modes. In addition to the nematoceran dipteran *Anopheles stephensi* (Cheatle-Jarvela et al., 2021), this progressive mode of segmentation has apparently also been observed in other holometabolous insect species including the lepidopteran *Bombyx mori* (the silk moth)(Nakao, 2010), the hymenopterans *Apis mellifera* (the honey bee)(Fleig, 1990; Binner and Sander, 1997) and *Nasonia vitripennis* (Rosenberg et al., 2014), and the coleopteran *Callosobruchus maculatus* (the cowpea weevil)(Patel et al., 1994).

In contrast, for short/intermediate germ-band insects, the full complement of segments is not defined prior to the onset of gastrulation (Patel et al., 1994) and some stripes of Engrailed/*wingless* will form prior to the completion of gastrulation, with other stripes forming after gastrulation has finished, in concert with the formation of additional segments from a posterior zone of growth (Akam, 1994; Choe et al., 2006; Davis & Patel, 2002; Strobl & Stelzer, 2014). Our results suggest that for the three ant species studied here, the striped patterns of expression of Engrailed and *wingless* appear after gastrulation. However, the stripes appear progressively, from anterior to posterior, after most of the growth of the embryo has occurred. At the molecular level, the fully elongated germ-band is therefore partitioned secondarily into segmental units. This is substantially different from what is seen in short/intermediate germ-band development, during which individual Engrailed strips appear within the posterior growth zone, in concert with the formation of new segments.

Accordingly, the results of our studies of *wg* mRNA and Engrailed protein expression described above indicate that the process of segmentation in these ant species does not occur via the simultaneous mode observed in the long germ-band-developing fly *D. melanogaster*, nor via the sequential mode seen in the short germ-band-developing locust *S. gregaria*. Rather, segmentation in these ants is more reminiscent of the progressive pattern of segmentation that has recently been described (Rosenberg et al., 2014, Cheatle-Jarvela et al., 2021). Curiously, this progressive mode of segmentation observed in the ants is coupled with an early pattern of embryogenesis reminiscent of short germ-band-developing genera. This mosaic combination of short and long germ-band patterns of embryogenesis, which to our knowledge seems not to have been described previously, adds to the diversity of patterns of insect embryogenesis that suggests the existence of a continuum of intermediates between the ancestral, short germ-band developing, sequentially-segmenting insects and the more evolutionarily derived long germ-band-developing, simultaneously-segmenting insects.

The simultaneous and progressive patterns of segmentation are considered to have arisen from the more ancestral sequential mode of segmentation as heterochronic shifts in the timing of segment formation (Davis & Patel, 2002; Clark et al., 2019). Indeed, a number of genes have been identified as “timing factors”, which may be responsible for these heterochronic shifts (Cheatle-Jarvela et al., 2021; Clark & Akam, 2016; Clark & Peal, 2018; Koromila et al., 2020; Rudolf et al., 2020; Soluri et al., 2020; Taylor & Deardon, 2021; Zhu et al., 2017). One of the obvious consequences of heterochronic changes in the patterns of insect embryogenesis is the differences in the time necessary for fertilized eggs to produce hatching larvae or nymphs in different species. It has recently been shown that there is little correlation between the size of insect eggs and their rate of development to adulthood (Church et al., 2019). Nevertheless, it is noteworthy that despite the similar sizes of their eggs and hatching larvae, the ant species studied here require from 10 to 20 days to progress from oviposition to hatching of first instar larvae, while the *Drosophila* embryo requires only a single day (Wieschaus & Nüsslein-Volhard, 1986). While the evolutionary constraints controlling the rate and duration of embryonic development remain poorly understood, it is likely that the trajectory, rates, and duration of embryonic development are constrained by overall life history and in particular by selective pressures exerted on the fertile egg-producing female. Although *Drosophila* females expend considerable effort in producing eggs that are well stocked with sufficient organelles and macromolecules to support rapid embryogenesis, they provide little parental care aside from avoidance of oviposition sites where parasitoids occur (Kacsoh et al., 2013; Ebrahim et al., 2015). Eggs are typically laid in rotting fruit where there can be intense competition for access to food by larvae, a situation which would select for rapid development of hatching larvae. As one adaptation to the rapid production and development of embryos, *Drosophila* eggs are produced in polytrophic meroistic ovaries, in which each developing oocyte is associated with a complex of 15 polyploid nurse cells that provide the majority of organelles and other synthetic machinery that will be required to support the development of the fertilized egg during embryogenesis. This rapid embryonic development is also supported by developmental control mechanisms involving significant maternal pre-patterning (St Johnston & Nüsslein-Volhard, 1992). In contrast to *Drosophila*, in the ants there is considerable care of eggs and embryos, either by the foundress queen or by workers in mature colonies. Because of this care in the relative absence of larval-larval competition, rapid embryonic development is less likely to be critical to the survival of ant larvae, compared to the solitary *Drosophila* developing under sometimes intense larval-larval competition.

Like *Drosophila* and the ant species studied here, the eggs of the hymenopteran species *Apis mellifera* (the honeybee), and *Nasonia vitripennis* (the jewel wasp) develop in polytrophic, meroistic ovaries. *Nasonia* embryos develop to hatching in 1-2 days while *Apis* embryos require three days to develop to hatching (Page & Peng, 2001; Zhang et al., 2019). In addition to the high morphological and taxonomic diversity in the Hymenoptera (Grimaldi & Engel, 2005; Aguiar et al., 2013), there exists great variation in life-history between hymenopteran lineages, and it is therefore likely that the different rates of embryogenesis observed across ants, bees, and wasps may have been selected as embryological adaptations contributing to life-history traits (e.g., life span) or other specializations (e.g., parasitism; solitary versus social life).

Insofar as the eggs of the ant species studied here are also produced in polytrophic, meroistic ovaries (Fig. 9), it is somewhat surprising that the ant embryos initially generate only small embryonic primordia and undergo a much longer period of embryogenesis than the other hymenopteran species noted here. Despite the strong synthetic capacity of polytrophic, meroistic ovaries, constraints are in place to direct a more lengthy period of embryogenesis than in these other hymenopteran species. One important distinction between *Apis mellifera* and *Nasonia vitripennis* compared to the ant species studied here is that the first brood of embryos produced by founder ant queens require those reproductive females both to provision their eggs with nutrition, macromolecules, and organelles necessary for embryogenesis and to care for and provide sustenance to their progeny after hatching ((Höldobler and Wilson, 1990). In contrast, reproductive females of *Nasonia* and *Apis* need only provision their eggs with nutrition that will carry the progeny through embryogenesis. *Nasonia* are parasitoids and lay their eggs within the pupae of carrion flies, a ready source of nutrients for hatched larvae. In the case of *Apis*, newly fertilized queens that found new hives do so together with a large escort of worker bees from their parental hive, who will care for the queen’s progeny larvae after they hatch. We suggest that this requirement of ant founder queens to provide both pre- and post-embryonic sustenance to their first brood places constraints on the size of the embryonic primordium and consequently, the rate of embryonic development.

**Figure 9.**
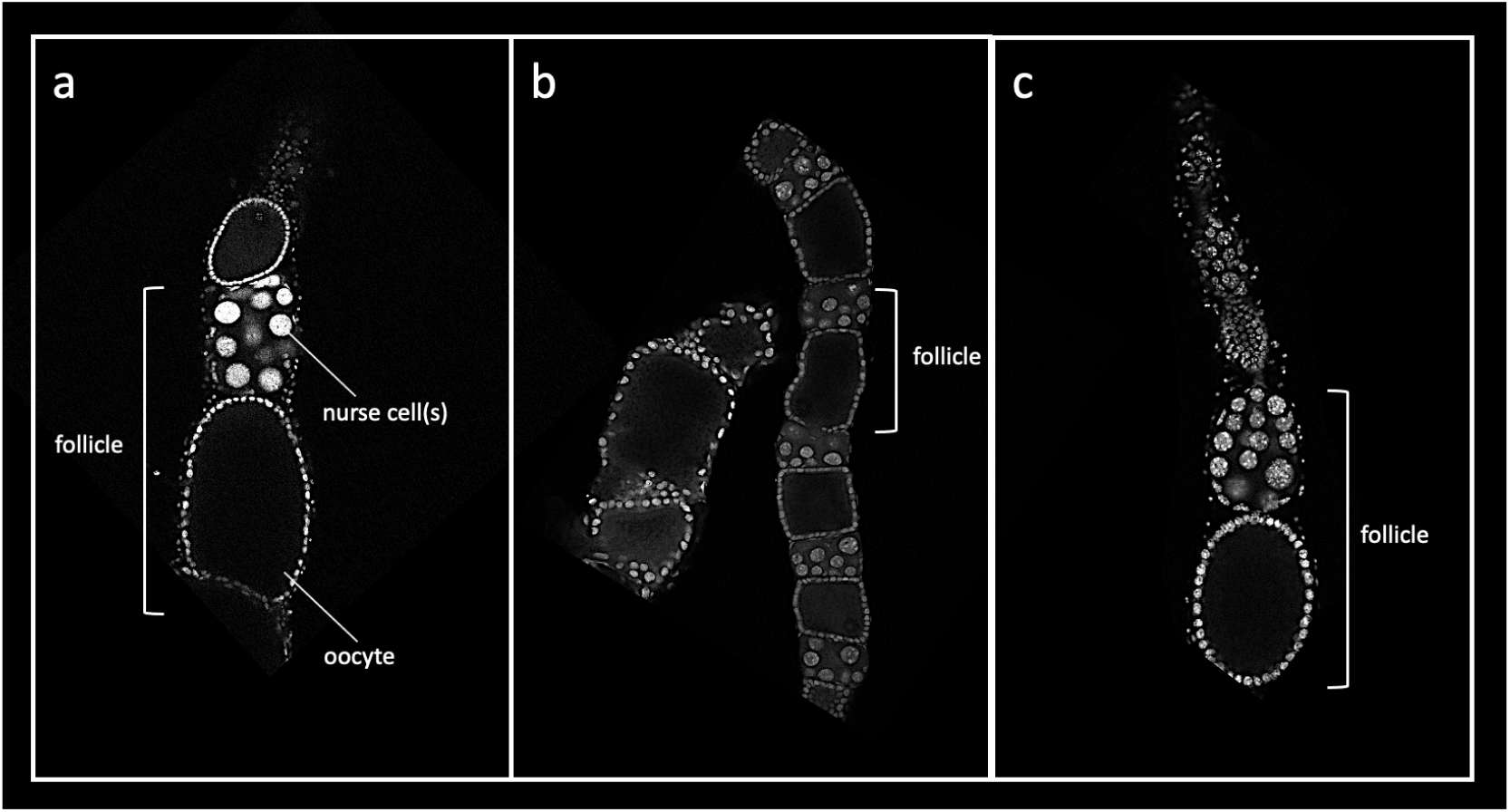
Comparison of ovarian follicles (egg chambers) from *Atta texana* (a), *Mycocepurus smithii* (b), and *Solenopsis invicta* (c). Each bracket indicates the position of a single egg chamber with the nurse cells positioned at top and the oocyte and associated follicle cells at bottom. Nuclei are stained with DAPI. The three myrmicine ants share the same polytrophic meroistic type of follicle, in which each oocyte obtains macromolecules and organelles from an associated cluster of nurse cells.

In eusocial insects, like ants, a critical period in the life cycle of the colony is the initial founding stage extending from dispersal of reproductive females to the appearance of the first brood of workers (Oster & Wilson, 1978). Recently, Fang et al. (2020) showed that for *M. smithii*, one of the species studied here, success in establishing a new colony by an individual queen is highly dependent upon the number of workers present in the newly forming colony. The presence of too few or too many *M. smithii* workers therefore decreases the likelihood of colony establishment, supporting the notion that intense selective pressure is also exerted upon the foundress queen in producing the optimal number of workers within this optimal time frame. Thus, mechanisms that ensure that the queen can nurture to adulthood a sufficient number of workers to establish the colony are likely to be important targets of selection. Moreover, a slow developmental rate may be optimal, particularly in species such as *A. texana* and *M. smithii,* for which foundress queens must divide their efforts and resources between producing and caring for brood and establishing and cultivating a nascent fungal garden during a time, at least in the case of *A. texana*, during which the queen does not forage for food. In this regard, the 1.5- to 2-fold increase in the duration of embryogenesis in the two fungus growing ant species, in comparison to *S. invicta*, whose queens are not tasked with cultivating a fungal garden, is also noteworthy. An additional constraint placed upon the level of nutritive investment that the queen can make in generating her eggs is the requirement for her to survive and remain fertile following the rigors of producing and tending her first brood, a constraint that is not placed upon females of insect species like *Drosophila*, which can produce a large number of eggs before mothers die at the end of a relatively short lifespan of about 50 days under optimal conditions (Linford et al., 2013).

The selective pressure applied upon the founder ant queen may also contribute to the introduction of developmental heterochrony within the life-time of the ant colony. This is consistent with the observation that the first clutch of worker progeny produced by the founder queen in *A. texana* take a shorter time to develop to adulthood than later cohorts do. The fitness strategy for a foundress queen is to maximize survival and brood production during a short period when workers are absent (Lotka, 1925; Marti et al., 2015). Foundress queens are therefore typically time-constrained (i.e., the queen needs to produce her first workers fast) and nutritionally constrained (i.e., the queen has limited resources in her body to produce the first brood of embryos and to feed and care for them after they hatch). For example, in the paper wasp, *Polistes fuscatus*, foundress queens reduce the size of first-emerging workers, which consequently developed faster to help in maintaining the colony (Reeve et al., 1998). Ant queens typically also produce smaller first (nanitic) workers (Espadaler & Rey, 2001; Porter & Tschinkel, 1986; Tschinkel, 1988), to conserve stored resources, and to increase the number of, and accelerate development of the first cohort of workers. Moreover, in the black garden ant, *Lasius niger*, the founder queen takes advantage of higher temperatures to accelerate development of the first brood (Kipyatkov et al., 2004). Because nanitic workers occur also in leafcutter ants (Wetterer, 1994; Wilson, 1980), we compared the timing of embryogenesis of *A. texana* foundress-laid eggs and mature-queen-laid eggs. Foundress-laid eggs needed only 75% of the total time to complete embryogenesis compared to eggs laid by mature queens (Fig. 6). Moreover, the smaller size of nanitic workers presumably requires fewer somatic cells, consequently decreasing the time needed to complete the elongation and segmentation stages. Goetsch (1937) (see also Metzl et al., 2018) observed that the first eggs laid and reared by *Pheidole pallidula* and queens produced minims, smaller than workers that would be produced by later broods. Interestingly, when Goetsch transferred the first eggs laid by founding queens to mature colonies in which the hatched larvae received nutrition from workers, they still developed into minim-sized workers, arguing that properties imbued by the founder queen into her first eggs determine their developmental destiny.

Interestingly, our analysis of the duration of embryogenesis in these ant species suggest that in addition to the introduction of heterochrony at the evolutionary scale in the development of modes of segmentation, heterochrony in embryogenesis can also be observed within an individual colony responding to different selective constraints over its life-span.

Although, as described above, the selective pressures that influence germ-band type, embryonic timing, and duration are likely to be diverse and related to life history, behavior and environment, it is clear that a variety of differences in the pattern of gene expression in the embryos affecting patterning, specification, growth, differentiation and morphogenesis must also underlie this diversity. Our analyses of the expression of Engrailed and *wingless* and the similarity of their final expression patterns with that of other insects support the designation of the extended germ-band stage as the phylotypic state in the insects (Domozet-Lošo & Tautz, 2010; Kalinka et al., 2010; Sander, 1983). However, within the insect lineage, and even among the hymenopteran species, there is considerable variation in the route through which embryos arrive at the extended germ-band stage, particularly in the earliest stages of embryogenesis. The differences between long, intermediate and short germ-band modes of development represent a portion of that variation. Embryogenesis in the ants that we have examined display a mosaic pattern with features of both short and long germband development. The embryonic primordium are short and require considerable growth in order to fill the volume of the egg. However, segments and segment polarity gene expression stripes do not appear one at a time, in tandem with growth. Rather, in the ants, segments and Engrailed/*wg* stripes appear at discrete positions along the extended anterior-posterior axis, as in *Drosophila*. It is tempting to speculate that the elaboration of segments and stripes in the ant species represents a transition between short/intermediate and long germ-band modes of development with the requirement for embryonic growth and the consequent longer duration in ant embryos making the progressive formation of segments/stripes from anterior-to-posterior of the axis more obvious and occurring later in embryogenesis than in *Drosophila*.

## Conclusion

Our observations suggest that evolutionary “tinkering” with the duration of processes, as well as the relative timing of distinct programs during embryogenesis may contribute considerably to the success of insects displaying different life history strategies. It is important to recognize that the differences between long, intermediate, and short germ-band development, and between sequential, progressive and simultaneous segmentation do not represent the full range of diversity in insect embryo developmental strategies. There exists considerable additional diversity in embryonic mechanisms of development, one example of which is the phenomenon of polyembryonic development displayed by the parasitoid wasp *Copidosoma floridanum* (Grbic et al., 1998). In this organism, the female lays a single egg into the egg of a host moth egg. Early development occurs within a cellular, rather than a syncytial environment, initially forming a primary morula. As development progresses, that primary morula subdivides, ultimately forming about 2000 proliferating morula, each of which is the primordium of a single embryo. Those primordia go on to undergo morphogenesis, germ-band extension and retraction, and formation of larvae. However, despite reaching the germ-band stage in a manner that is dramatically different, these embryos exhibit a pattern of Engrailed stripes along the anterior/posterior axis that is very similar to that ultimately observed in the ants (Grbic et al., 1996, 1998). This exceptional polyembryonic example, together with our observations that the patterns of segment gene expression are reached in ants via a process somewhat different from that seen in representative short, intermediate, and long germ-band-developing insects, as well as the newly recognized progressive mode, supports the existence of tremendous plasticity in the nature and trajectory of developmental events employed to reach the phylotypic state. Our studies represent an early step in better understanding the details of the process of embryogenesis and of how their unique life history traits influence that process in the ants.

## Acknowledgements

We thank Etzel Garcia, Peter Duong, James Kurian, and Ryan Bailey for colony maintenance; Paul Macdonald for experimental suggestions and assistance with microscopy. The study was funded by a Texas Ecolab award and a UT Austin Global Research Fellowship to CCF, National Science Foundation award DEB-1354666 to UGM, and the W.M. Wheeler Lost Pines Endowment from the University of Texas at Austin. This work was also supported by a Discovery Grant from the Natural Sciences and Engineering Research Council of Canada (NSERC) to E.A. and doctoral fellowship from FRQNT (Quebec) to A.R.

## Appendices

### Appendix A

**Table B1.**
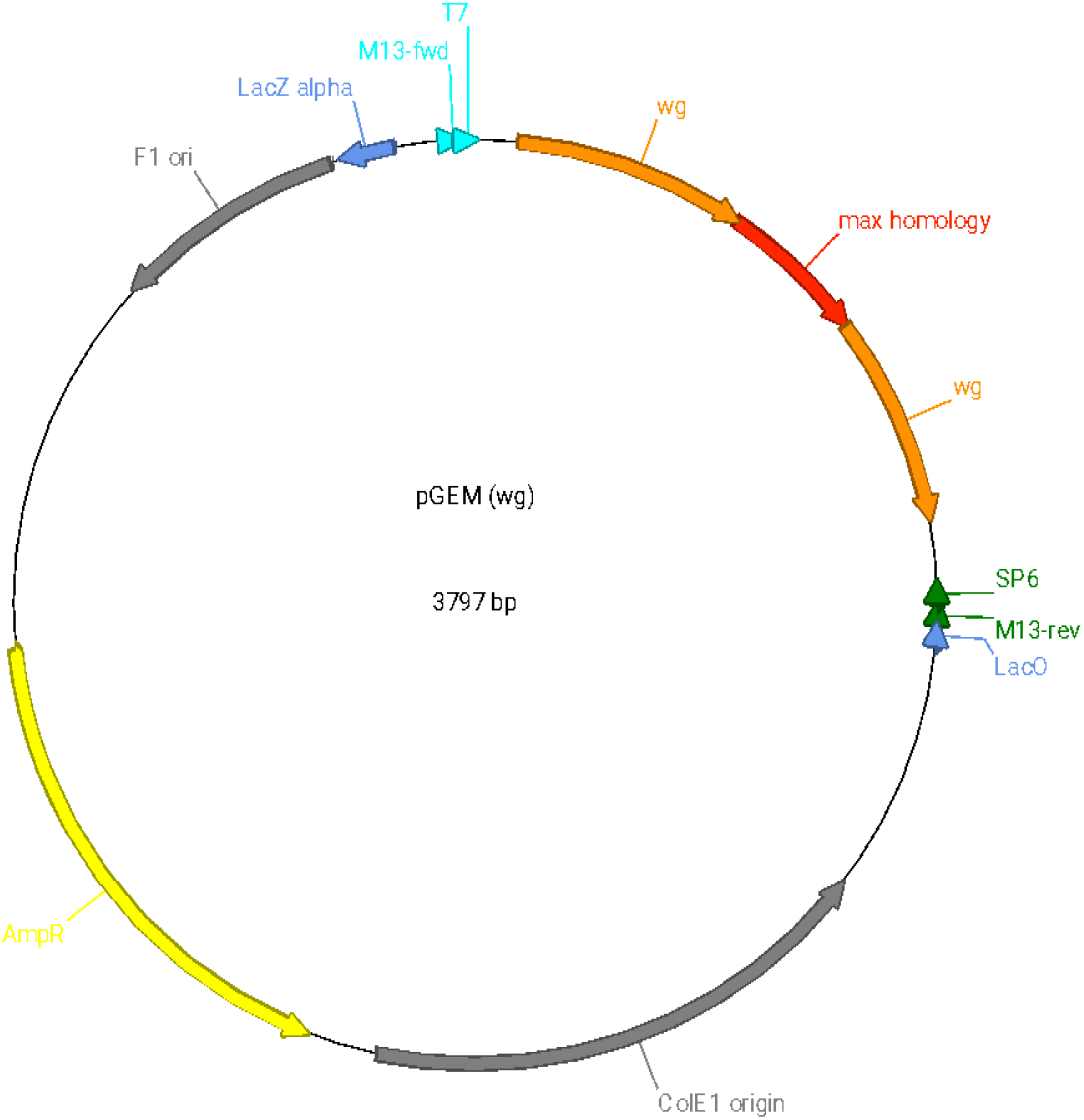
Plasmid for amplifying *Atta colombica* Wnt-1 mRNA. We designed the *wingless* probe based on the sequence of that gene in the published genome of *Atta colombica* (Wnt-1 mRNA; NCBI reference sequence: XM_018204929.1). The sequence of wingless mRNA (orange and red regions) is listed below.

